# GMHI-webtool: a user-friendly browser application for assessing health through metagenomic gut microbiome profiling

**DOI:** 10.1101/2022.06.30.498296

**Authors:** Daniel Chang, Vinod K. Gupta, Benjamin Hur, Kevin Y. Cunningham, Jaeyun Sung

## Abstract

**Summary:** We recently introduced the Gut Microbiome Health Index (GMHI), a stool-based indicator for monitoring health given the state of one’s gut microbiome. GMHI depends on health-prevalent and health-scarce species determined and validated using a pooled dataset of 5,026 stool shotgun metagenomic samples from 43 independent studies. Encouragingly, GMHI has already been utilized in various studies focusing on identifying differences in the gut microbiome between cases and controls. However, current computational barriers and logistical issues prevent researchers from computing, interpreting, and contextualizing GMHI, thereby limiting its further widespread utilization. Herein, we introduce the GMHI-webtool, a user-friendly browser application that computes GMHI, health-prevalent/scarce species, α-diversities, and taxonomic distributions of the gut microbiome from stool samples. Users of our interactive online tool can visualize their results and compare side-by-side with those from our pooled reference dataset, as well as export data in .csv format and high-resolution figures.

**Availability and implementation:** GMHI-webtool is freely available here: https://gmhi-webtool.github.io/. Source code: https://github.com/danielchang2002/GMHI-webtool.

## Introduction

To date, human gut microbiome research has given us various convincing associations and mechanistic insights into chronic diseases (Duvallet et al. 2017; Fan and Pedersen 2020; Miyauchi et al. 2022; J. Sung et al. 2017), as well as promising predictive tools for clinical applications (V. K. Gupta et al. 2021; Ananthakrishnan et al. 2017). To demonstrate the utility of gut microbiome data for public health applications, we have recently introduced the Gut Microbiome Health Index (GMHI), a stool metagenome-based indicator for monitoring health (Vinod K. Gupta et al. 2020). GMHI is a biologically interpretable mathematical formula for predicting the likelihood of disease independent of the clinical diagnosis. In brief, GMHI was computed from two sets of microbial species associated with healthy and disease conditions (i.e., “health-prevalent” and “health-scarce” species, respectively), and was determined and validated using a pooled dataset of 5,026 stool shotgun metagenomic samples from 43 independent studies. As a proof-of-concept, our index achieved a balanced accuracy of 73.7% in predicting whether a person had a clinically diagnosed disease (or not) on an external validation set of stool metagenomes from 679 people, outperforming methods based solely upon ecological indices and a random forest classifier.

Since its original publication in 2020, GMHI has been used in studies focused on the effects of environmental (Gacesa et al. 2022), and genetic/socioeconomic (Xu et al. 2022) factors on the human gut microbiome and health. In these studies, GMHI was computed on stool metagenomes to compare groups of interest, such as different age cohorts, case/controls, and medication users/non-users. Moreover, health-prevalent/scarce species were used to confirm the health relevance of newly computed species sets, such as the “Longevous Gut Microbiota Signature” species set found by Xu *et al*. As an interesting extension of our tool to household pets, the GMHI strategy was applied to assess the health of cats (C.-H. Sung et al. 2022).

Despite the promising utility of GMHI as a noninvasive and dynamic tool to assess gut health, a few logistical issues can be addressed to improve its widespread applicability. While routine data pre-processing, such as taxonomic profiling using MetaPhlAn2 (Truong et al. 2015), can be done by researchers who are knowledgeable of bioinformatics tools and basic programming, novel downstream analyses may need to be directed by those who are relatively unfamiliar with handling computational pipelines. In the case of computing GMHI, proficiency in R programming and its external libraries are required. Similarly, meticulous text parsing is required to identify the presence of health-prevalent/scarce species, a task vital to GMHI’s biological interpretability. Moreover, because databases of GMHI scores are not yet available, it is thus infeasible to contextualize GMHI scores by comparing them to a reference control population. We posit that these challenges must be overcome for the widespread adoption and implementation of GMHI in clinical settings.

Here, we present the GMHI-webtool, a user-friendly browser application that allows anyone with interest in GMHI to compute the index and various other ecological indices from a stool metagenome sample. Specifically, after performing MetaPhlAn2 on a metagenome and uploading the output taxonomy profile, users can extract the following: (i) GMHI; (ii) health-prevalent and health-scarce species detected in the sample; (iii) several α-diversity indices: richness, evenness, Shannon, inverse Simpson; and (iv) distribution of relative abundances of various taxonomies. Additionally, users can visualize and compare these results with healthy (i.e., self-reported to not having a disease or disease-related symptoms) and nonhealthy (i.e., patients with 1 of 12 different disease or abnormal body-weight conditions) populations from a pooled reference gut microbiome dataset of 5,026 stool shotgun metagenomic samples, as well as export data (.csv) and high-resolution figures (.png or .svg).

## Implementation

GMHI-webtool is a client-side JavaScript application written using the D3.js (Bostock, Ogievetsky, and Heer 2011) library. Python and the scikit-learn (Pedregosa, Varoquaux, and Gramfort, n.d.) library were used to pre-compute GMHI, α-diversity indices, and Principal Component Analysis (PCA) of the pooled dataset of 5,026 stool shotgun metagenomic samples. See **Supplementary Information** for a detailed description of the software used.

## Usage

Before using the GMHI-webtool, users need to upload the unedited .txt taxonomic profile(s) from running MetaPhlAn2 on metagenome .fastq/.fastq.gz/.bam files(s) (see **Supplementary Information** for full pipeline). GMHI-webtool supports both single-sample and multi-sample versions of the MetaPhlAn2 output. Users can select whether to compare side-by-side the input sample with healthy or nonhealthy (or both) populations from our aforementioned pooled dataset (**Supplementary Fig. S1**).

GMHI of the input sample, along with α-diversity indices (richness, evenness, Shannon diversity, and inverse Simpson diversity) (see **Supplementary Information** for details), are computed and shown in relation to those of samples from the pooled dataset (**Fig. 1A**). Users can export these values as a .csv file by clicking the “Export as CSV” button.

**Fig. 1.**
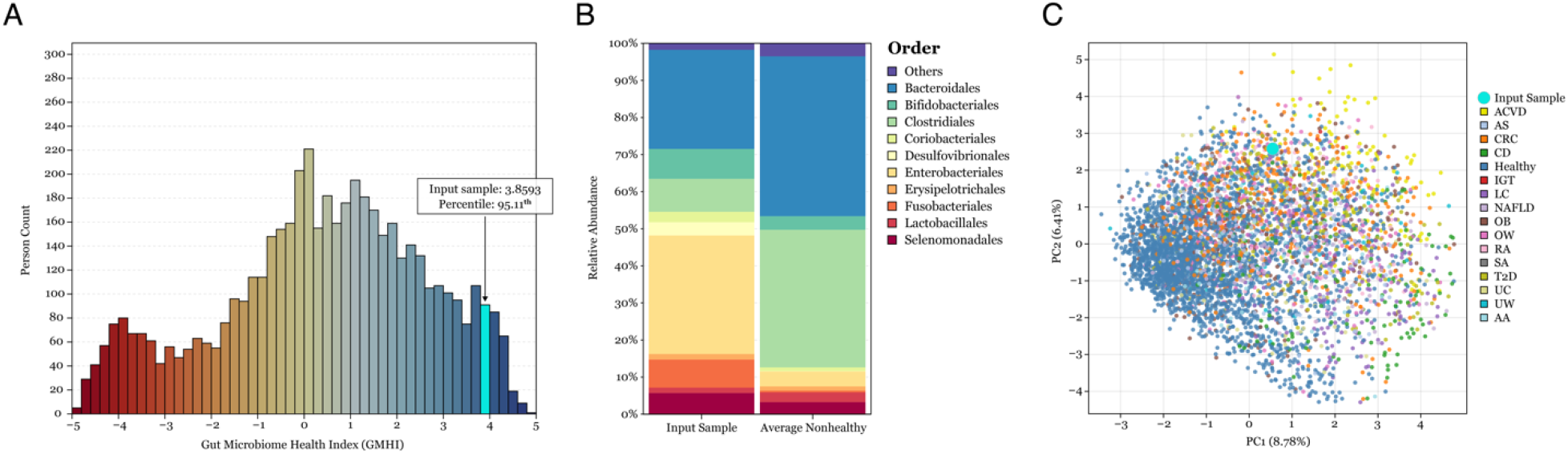
Example visualizations from GMHI-webtool using an example gut microbiome (stool metagenome) sample. **(A)** GMHI of an input sample (cyan) in relation to all 5,026 gut microbiome metagenome samples in our pooled reference dataset. **(B)** The relative abundances of microbial organisms (at the order level) in an input sample (left). Averages across nonhealthy samples (right). **(C)** A PCA plot showing the projection of an input sample (cyan) amongst microbiome samples of our pooled dataset.

Additionally, our webtool presents stacked bar charts describing the distribution of taxa relative abundances of the input sample at the phylum, class, order, and family level. Average relative abundances of any of the two populations in our pooled dataset are also shown (**Fig. 1B**). Users can hover the mouse over a taxon name or its corresponding bar to view its relative abundance (**Supplementary Fig. S2**).

A PCA plot of microbiome samples from our pooled dataset, along with the option to project the user’s input sample onto the ordination plot, is available (**Fig. 1C**). Users can hover the mouse over phenotype acronyms to read the full phenotype name and observe the input sample’s projected position in relation to different phenotypes.

A table describing the relative abundances of health-prevalent/scarce species in the input sample, along with their median values in the healthy and nonhealthy populations in our pooled dataset, is available (**Supplementary Fig. S3**). Likewise, a table of the top three most abundant taxa of each taxonomic rank, including comparisons with the pooled dataset populations, is available (**Supplementary Fig. S4**).

## Conclusion

GMHI-webtool is a browser application with an intuitive and simple user interface that allows researchers to easily analyze GMHI, α-diversity indices, and taxonomic distributions of a stool metagenome sample. Importantly, the aim of our webtool is to democratize the ability to gain important health insights from gut microbiome data. By addressing the current computational barriers, our GMHI-webtool can facilitate future clinical applications of gut microbiome research.

## Acknowledgment

This work was supported in part by the Mayo Clinic Center for Individualized Medicine and a Translational Product Development Fund Program Award from the Minnesota Partnership for Biotechnology and Medical Genomics (to J.S.).

