## Supplementary Information for "GMHI-webtool: a user-friendly browser application for assessing health through metagenomic gut microbiome profiling"

for

Daniel Chang *et al.*

4 References 9

### 1 Webtool Information

#### 1.1 Implementation

Python and the Numpy (Harris et al., 2020) library were used to pre-compute GMHI, $\alpha$-diversities, and taxonomic distributions of the pooled dataset. The scikit-learn (Pedregosa et al., 2011) library was used for Principal Component Analysis (PCA) of the pooled dataset. Access the data here. Regarding the front-end, GMHI-webtool is a client-side application written using JavaScript and the D3.js (Bostock, 2012) library. The Math.js (de Jong and Mansfield, 2018) library is used to project the input sample onto the first two principal components of the pooled dataset. $\alpha$-diversities of the input sample, along with text parsing and validation, are implemented using JavaScript functions.

#### 1.2 User Input

Users need to first upload (**Supplementary Fig. S1A**) the taxonomy profile (see **Section 2**). If the file has multiple samples, users can select the sample for analysis (**Supplementary Fig. S1B**). Users may choose to compare the input sample with healthy samples, nonhealthy samples, or all samples in the pooled gut microbiome dataset (**Supplementary Fig. S1C**). This selection pertains to the first two plots only. After making the proper selections, users can press the "display results" button (**Supplementary Fig. S1D**) to compute and visualize the analyses.

#### 1.3 User Interaction and Exports

Users can change plot options (e.g., index, taxonomic level) by using the tabs above plots (**Supplementary Fig. S2A**). Additionally, users can hover their mouse over legend text (**Supplementary Fig. S2B**) to highlight information.

Users can export index data by clicking the "export as CSV" button (**Supplementary Fig. S1E**), and export plots by clicking links below them (**Supplementary Fig. S2C**).

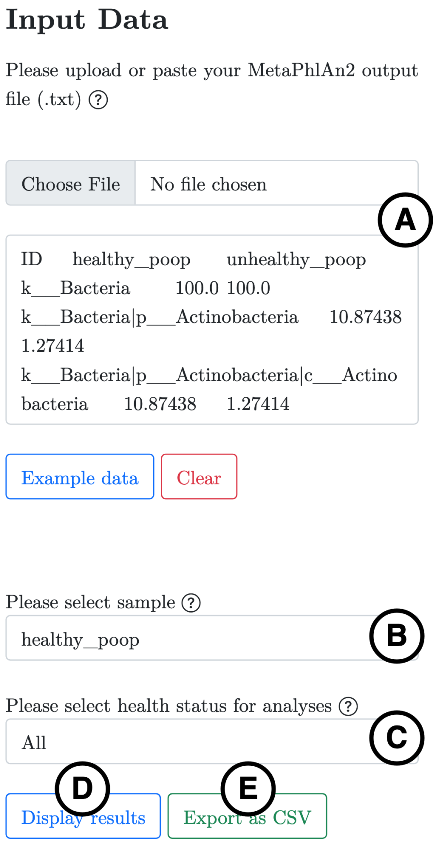

**Supplementary Fig. S1.** The user input panel.

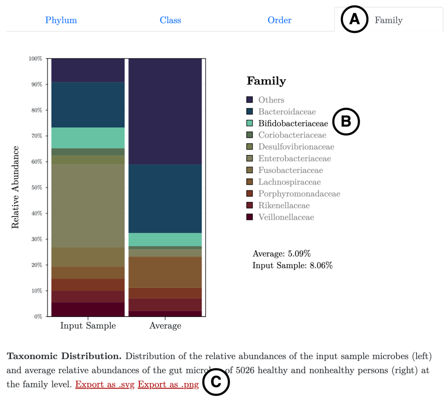

**Supplementary Fig. S2.** Stacked bar plots of taxonomic distributions.

#### 1.4 Data Tables

GMHI-webtool provides a table describing the relative abundances of health-prevalent/scarce species in the input sample, along with their median values in the healthy and nonhealthy populations in our pooled dataset **(Supplementary Fig. S3)**. Likewise, a table of the top three most abundant taxa of each taxonomic rank, including comparisons with the pooled dataset populations, is available **(Supplementary Fig. S4)**.

Health-prevalent and Health-scarce Species

| Species Name | Presence | Relative Abundance | Median (Healthy) | Median (Nonhealthy) |
|---|---|---|---|---|
| *Alistipes senegalensis* | ✅ | 0.086% | 0.004% | 0.000% |
| *Bacteroidales bacterium_ph8* | ✅ | 0.093% | 0.128% | 0.000% |
| *Bifidobacterium adolescentis* | ✅ | 4.700% | 0.291% | 0.000% |
| *Bifidobacterium angulatum* | ❌ | - | 0.000% | 0.000% |
| *Bifidobacterium catenulatum* | ✅ | 0.400% | 0.000% | 0.000% |
| *Lachnospiraceae bacterium_8_1_57FAA* | ❌ | - | 0.000% | 0.000% |
| *Sutterella wadsworthensis* | ✅ | 1.598% | 0.000% | 0.000% |
| *Anaerotruncus colihominis* | ❌ | - | 0.000% | 0.000% |
| *Atopobium parvulum* | ❌ | - | 0.000% | 0.000% |
| *Bifidobacterium dentium* | ❌ | - | 0.000% | 0.000% |
| *Blautia producta* | ✅ | 0.026% | 0.000% | 0.000% |
| *candidate division_TM7_single_cell_isolate_TM7c* | ❌ | - | 0.000% | 0.000% |
| *Clostridiales bacterium_1_7_47FAA* | ✅ | 0.036% | 0.000% | 0.000% |
| *Clostridium asparagiforme* | ✅ | 0.012% | 0.000% | 0.000% |
| *Clostridium bolteae* | ✅ | 0.010% | 0.000% | 0.018% |
| *Clostridium citroniae* | ✅ | 0.222% | 0.000% | 0.000% |
| *Clostridium clostridioforme* | ✅ | 0.003% | 0.000% | 0.000% |
| *Clostridium hathewayi* | ✅ | 0.108% | 0.000% | 0.003% |
| *Clostridium nexile* | ✅ | 0.059% | 0.000% | 0.000% |

**Supplementary Fig. S3.** Table of health-prevalent and health-scarce species.

Most Abundant Taxa

| Rank | Most Abundant Taxa | Relative Abundance | Median in Healthy | Median in Nonhealthy | Median in All |
|---|---|---|---|---|---|
| **Phylum** | Proteobacteria | 29.416% | 1.104% | 1.815% | 1.322% |
| | Firmicutes | 24.886% | 47.080% | 45.954% | 46.657% |
| | Bacteroidetes | 21.133% | 38.169% | 39.335% | 38.829% |
| **Class** | Gammaproteobacteria | 22.540% | 0.199% | 0.737% | 0.331% |
| | Bacteroidia | 21.133% | 38.169% | 39.335% | 38.829% |
| | Actinobacteria | 16.040% | 2.803% | 1.401% | 2.001% |
| **Order** | Enterobacteriales | 22.538% | 0.098% | 0.484% | 0.195% |
| | Bacteroidales | 21.133% | 38.169% | 39.335% | 38.829% |
| | Bifidobacteriales | 15.481% | 1.809% | 0.612% | 1.177% |
| **Family** | Enterobacteriaceae | 22.538% | 0.098% | 0.484% | 0.195% |
| | Bifidobacteriaceae | 15.481% | 1.809% | 0.612% | 1.177% |
| | Bacteroidaceae | 13.917% | 15.781% | 20.829% | 17.452% |
| **Genus** | Escherichia | 21.517% | 0.077% | 0.287% | 0.137% |
| | Bifidobacterium | 15.481% | 1.809% | 0.608% | 1.168% |
| | Bacteroides | 13.917% | 15.781% | 20.829% | 17.452% |
| **Species** | Escherichia coli | 16.809% | 0.043% | 0.226% | 0.092% |
| | Fusobacterium varium | 8.054% | 0.000% | 0.000% | 0.000% |
| | Bifidobacterium longum | 6.941% | 0.333% | 0.121% | 0.223% |

**Supplementary Fig. S4.** Table of the top three most abundant taxa of each taxonomic rank.

### 2 Pipeline

Prior to using GMHI-webtool, users need to profile their metagenome .fastq files using MetaPhlAn2 (Truong et al., 2015). The following is the pipeline used to preprocess and profile the stool metagenomes used to compute GMHI (Gupta et al., 2020). However, users are free to use any other pipeline provided that taxonomic profiling uses MetaPhlAn2. For brevity, this pipeline assumes that paired-end metagenomes are used.

#### 2.1 Setup

Install/update softwares:

- bbmap (repair.sh) v38.93 (Bushnell, 2014)
- fastqc v0.11.9 [Source]
- bowtie2 v2.4.4 (Langmead and Salzberg, 2012)
- samtools v0.1.19 (Li et al., 2009)
- bedtools v2.30.0 (Quinlan and Hall, 2010)
- trimmomatic v0.39 (Bolger et al., 2014)
- MetaPhlAn2 v2.x.x (Truong et al., 2015)

```
# make sure the paired end metagenome files are available
ls
.
├── in1.fastq
└── in2.fastq

# set this var to directory containing human reference genome (use GRCh38/hg38)
# example:
$HUMAN_REFERENCE_GENOME=/users/mynamejeff/human_genomes/GRCh38_noalt_as/

# set this var to desired number of parallel processes
# example:
$N_JOBS=16

# set this var to directory containing metaphlan2 clade markers
# example:
$CLADE_MARKERS=/users/mynamejeff/metaphlan2_data/clade_markers

# Prepare adapter sequence file
```

```
echo ">PrefixPE/1
TACACTCTTTCCCTACACGACGCTCTTCCGATCT
>PrefixPE/2
GTGACTGGAGTTCAGACGTGTGCTCTTCCGATCT" > TruSeq3-PE.fa
```

##### 2.2 Repair fastq files using bbmap

```
repair.sh in1=in1.fastq in2=in2.fastq out1=repaired1.fastq \
out2=repaired2.fastq outs=garbage
```

##### 2.3 Quality check and identification of overrepresented sequences

```
fastqc repaired1.fastq
fastqc repaired2.fastq
unzip repaired1_fastq.zip
unzip repaired2_fastq.zip
```

##### 2.4 Extract probable adapter sequences from FastQC outputs

```
for f in repaired1_fastqc/fastqc_data.txt; do
    echo $f `grep -A100 ">>Overrepresented sequences" $f | \
    grep -m1 -B100 ">>END_MODULE" | \
    grep -P "Adapter|PCR" | \
    awk '{print ">overrepresented_sequences" "_" ++c "/1" $1}'` | \
    awk '{gsub(/\/1/,"/1\n")}1' | \
    awk '{gsub(/>/,"\n>")}1' | \
    awk '{gsub(/fastqc_data.txt/,"")}1' | \
    awk 'NF > 0';
done > adapter1.txt

for f in repaired2_fastqc/fastqc_data.txt; do
    echo $f `grep -A100 ">>Overrepresented sequences" $f | \
    grep -m1 -B100 ">>END_MODULE" | \
    grep -P "Adapter|PCR" | \
    awk '{print ">overrepresented_sequences" "_" ++c "/1" $1}'` | \
    awk '{gsub(/\/1/,"/1\n")}1' | \
    awk '{gsub(/>/,"\n>")}1' | \
    awk '{gsub(/fastqc_data.txt/,"")}1' | \
    awk 'NF > 0';
done > adapter2.txt
```

#### 2.5 Remove human contaminants

```
bowtie2 -p $N_JOBS -x $HUMAN_REFERENCE_GENOME -1 repaired1.fastq \
-2 repaired2.fastq -S mapped.sam

samtools view -bS mapped.sam > mapped.bam

samtools view -b -f 12 -F 256 mapped.bam > human.bam

samtools sort -n human.bam human_sorted -@ $N_JOBS

bedtools bamtofastq -i human_sorted.bam -fq human1.fastq -fq2 human2.fastq
```

#### 2.6 Remove adapter sequences and low quality reads

```
cat adapter1.txt adapter2.txt TruSeq3-PE.fa > adapters.txt

trimmomatic PE -threads $N_JOBS human1.fastq human2.fastq -baseout QC.fastq.gz \
ILLUMINACLIP:adapters.txt:2:30:10:2:keepBothReads LEADING:3 TRAILING:3 MINLEN:60
```

#### 2.7 Profile the metagenome

```
metaphlan2.py QC_1P.fastq.gz,QC_2P.fastq.gz --bowtie2db $CLADE_MARKERS \
--bowtie2out --index mpa_v20_m200 --nproc $N_JOBS --input_type fastq \
-o profiled_metagenome.txt
```

After running the pipeline, users can upload the taxonomic profile "profiled_metagenome.txt" to GMHI-webtool. Users can also use MetaPhlAn2 to merge multiple taxonomic profiles:

```
ls
.
├── profiled_metagenome1.txt
├── profiled_metagenome2.txt
└── profiled_metagenome3.txt

merge_metaphlan_tables.py profiled_metagenome*.txt > merged_abundance_table.txt
```

And upload the merged file "merged_abundance_table.txt" to GMHI-webtool.

### 3 $\alpha$-diversity Indices

GMHI-webtool computes a number of $\alpha$-diversities from stool metagenome samples. Let $p_i$ be the relative abundance of the $i$th species in the sample. For consistency with the original GMHI work, only the 313 species considered during the computation of GMHI were considered (Gupta et al., 2020). Let $S = 313$ be the maximum number of species in a single sample. Let $c = 0.00001$ be the presence threshold.

#### 3.1 Richness

Richness R is the number of species with relative abundance greater than the presence threshold.

$$R = |\{\, i \mid p_i > c \,\}|$$

#### 3.2 Shannon Diversity

Shannon Diversity $H'$ is derived from Shannon entropy, and is a measure of the uncertainty associated with predicting the species of any microbe in the sample.

$$H' = -\sum_{\forall i[p_i > 0]} p_i ln(p_i)$$

#### 3.3 Evenness

Evenness $E$ is a measure of close in number (or relative abundance) different species are.

$$E = \frac{H'}{ln(S)}$$

#### 3.4 Simpson Diversity

Simpson diversity is equivalent to the probability that two randomly selected microbes are of the same species (SIMPSON, 1949).

$$\lambda = \sum_{\forall i[p_i > 0]} ln(p_i)$$

#### 3.5 Inverse Simpson

Inverse Simpson diversity is the reciprocal of Simpson diversity.

$$I = \frac{1}{\lambda}$$
